# A Multi-Purpose Worm Tracker Based on FIM

**DOI:** 10.1101/352948

**Authors:** Matthias Kiel, Dimitri Berh, Jens Daniel, Nils Otto, Adrian ter Steege, Xiaoyi Jiang, Eva Liebau, Benjamin Risse

## Abstract

The analysis of behavioural traits of *Caenorhabditis elegans* is an important method for understanding neuromuscular functions and diseases. Since *C. elegans* is a small and translucent animal which conducts a variety of complex movement patterns many different imaging and tracking protocols are used for different behavioural traits. Thus a unified multi-purpose imaging and tracking system for multiple behavioural assays would be favourable to improve statistical strength and comparability. Here we present a novel worm tracking toolbox based on the FIM (<u>F</u>rustrated total internal reflection (FTIR) based <u>I</u>maging <u>M</u>ethod) system incorporating a variety of different behavioural assays into a single imaging and tracking setup.

First, we apply the FTIR-based imaging method to *C. elegans*, thus we are able to improve the overall image quality compared to state of the art recording techniques. This method is easy to use and can be utilised to image animals during crawling on agar and trashing in water. Second, we extended the existing FIMTrack software to extract skeleton-based posture and motion features of multiple worms with very high accuracy in a comparatively large field-of-view. Third, we integrated a variety of different assays into this system. We carried out chemotaxis assays both with attractant and repellent chemicals. A novel electrotaxis dome compatible with FIM allows locomotion analyses that are not corrupted by random aberrations in unrestricted movement. Additionally, the FIM based worm tracker is able to analyse thrashing behaviour of multiple worms automatically with a high accuracy. Finally we demonstrate the capacity of the FIM based worm tracker to observe GFP signals in *C. elegans* worms. We tested our new *C. elegans* tracking suite with mutant strains of the ubiquitin-fold modifier 1 (Ufm1) cascade. We identified intermediate chemosensory phenotypes in Ufm1 cascade mutants which were previously undetected.

## Introduction

*C. elegans* is an important model organism for studying neuronal networks which govern behaviours like movement and learning (Ardiel and Rankin 2010) since it offers several advantages like eutely, a short generation time and its translucent body so that internal organs can easily be observed using a microscope. However, whole body imaging to study its behaviour suffers from its small size and transparent appearance, especially since the recording has to be done on moist surfaces like agar or in water. Several different imaging and tracking systems exist up to now, many of them specialised towards a certain behavioural protocol (reviewed by (Husson et al. 2012)). In general, worm trackers can be distinguished into two types, single worm trackers and multi worm trackers. Single worm trackers offer a high resolution per worm and are able to extract detailed movement patterns. One of the first *C. elegans* trackers was able to deduce up to 144 different parameters (Feng et al. 2004). The Nemo tracking application has an implemented GUI to be more user-friendly and is also able to calculate various movement features (Tsibidis and Tavernarakis 2007). “Track-a-Worm” was developed to be easily implemented in other labs and includes a GUI, a motorised stage to follow the worm movement and many analysis modules (Wang and Wang 2013). Some trackers are specialised for specific phenotypic analysis like food ranging behaviour (Moy et al. 2015) or tracking worms in a 3D environment (Kwon et al. 2013). Other worm trackers specialize on swimming gaits and cannot be adapted to crawling behaviour (Krajacic et al. 2012; Restif and Metaxas 2008). Nevertheless, a shared disadvantage of single-worm trackers is their low throughput. Multi-worm trackers tackle this problem. The “Parallel Worm Tracker” can track several worms at a time. However a centroid based algorithm is used, resulting in a single point representation of the tracked animals (Ramot et al. 2008). Based on this representation the calculated parameters are restricted to basic features like the crawling path or speed. Therefore centroid-based systems are unsuitable for studying complex movement patterns like thrashing or omega bends. Yu and his colleges developed a specialised tracker to analyse activity of up to 12 worms in separate wells over a period of days to investigate development and mating behaviour (Yu et al. 2014). The broadly used “Multi-Worm Tracker” (MWT) incorporated the skeleton-based system, uses a high-resolution and can process a huge amount of data in real-time mode by a LabView based platform, where the imaging data itself is not stored (Swierczek et al. 2011). The MWT uses different parameters and thresholds to automatically reject all nonworms, obscured worms and touching worms from the analysis. This processing of the data leads to an increased analysis speed, but also to a possible loss of informative data, especially because a post processing of the data is not possible in real-time mode.

A common drawback of most worm trackers is the use of a bright-field illumination. Light reflections caused by the underground medium can create a high number of artefacts. In addition some behaviour experiments like thrashing rely on imaging worms under water, which can lead to further artefacts. To cope with illumination artefacts many trackers apply Gaussian filtering algorithms to homogenise the images (Moy et al. 2015) and convert the grey-scale image into a binary image to extract the animals from the background (Feng et al. 2004; Kwon et al. 2013; Tsibidis and Tavernarakis 2007; Wang and Wang 2013). However some applications require higher contrast images for faithful segmentation. Furthermore, devices for environmental stimulation which cover the agar plate would obscure the direct illumination. The system presented here addresses the above mentioned challenges directly by the imaging hardware and by image processing steps offering a broad flexibility in the overall assay design and is therefore an optimal tool to track and analyse small transparent *C. elegans.*

Our setup is an advancement of the FIM setup (FIM = <u>F</u>rustrated total internal reflection-based <u>I</u>maging <u>M</u>ethod (Risse et al. 2013)). FIM was first designed to visualize subtle changes in the locomotion patterns of *Drosophila* larvae (Risse et al. 2014a; Risse et al. 2014b). However, due to its high power in image acquisition, it turned out to be a powerful tool to image and quantify *C. elegans* as well. Instead of direct illumination, the FIM setup utilizes frustrated total internal reflection (FTIR) to illuminate the worms with infrared light. In consequence, illumination problems causing a low contrast between worms and background and possible false positive artefacts are reduced on the physical level. Due to this high contrast, thresholding is sufficient to extract the animals (i.e. foreground) faithfully from the images (Risse et al. 2013). Further analysis is done by using FIMTrack, a custom tailored associated software for the FIM setup, which is developed for automated behaviour analysis. Furthermore, FIMTrack offers the possibility to quickly remove “ghost trajectories” (e.g. dust particles), where ghost trajectories mean all trajectories shorter than a user defined frame number or outside a user specified region of interest. FIMTrack is freely available at fim.uni-muenster.de.

Most importantly, this system is capable of screening worm strains in different behavioural paradigms like chemotaxis, electrotaxis and thrashing with high throughput. Furthermore, the use of GFP-labelled *C. elegans* is possible and could facilitate a greater range of experiments. In this study we demonstrate the reliability and suitability of our imaging and tracking system in the analysis of even subtle phenotypic differences observed in ubiquitin-fold modifier 1 (Ufm1)cascade mutant strains in multiple behavioural experiments. Ufm1 is a posttranslational modification system that belongs to the ubiquitin-like protein family. Maturation and deconjugation of Ufm1 is catalyzed by the Ufm1-specific protease, mature Ufm1 is then activated by Uba5 (Ufm1 activating enzyme 5, E1), conjugated by Ufc1 (Ufm1 conjugation enzyme 1, E2) and ligated by Ufl1 (Ufm1 ligating enzyme 1, E3). While the cascade has been shown to be involved in cellular homeostasis, influencing cell division, growth and endoplasmic reticulum functions, its biological role is still poorly understood (Daniel and Liebau 2014). Additionally, a study by Chen et al. revealed that the loss of Ufm1-specific protease (Ufsp2) leads to a reduced chemosensory reaction, whereas other Ufm1 cascade proteins, like Ufc1 and Uba5, showed no altered chemotaxis response (Chen et al. 2014). As part of our extensive phenotyping of the respective deletion mutants, we investigated the role of the Ufm1 cascade in different behavior experiments using the novel FIM tracking approach in order to reproduce the chemosensory defect of Δ*ufsp-2* mutants as well as to demonstrate the suitability of our novel tracking system. Due to the very high precision of our system we even detected intermediate phenotypes in other Ufm1 cascade mutants.

## Methods

### Worm Strains and Culturing

The *C. elegans* strains wild type (Bristol N2) and the deletion mutants Δ*ufc-1* (tm4888), Δ*uba-5* (ok3364), Δ*ufsp-2* (tm5790) and Δ*odr-3* (n1605) were obtained from the Caenorhabditis Genome Center (CGC), University of Minnesota. Worms were cultured at 20 °C on standard NGM plates seeded with the *Escherichia coli* OP50 strain following standard protocols (Brenner 1974). All behaviour experiments were conducted with synchronised young adult worms. Synchronisation of *C. elegans* strains were performed by the hypochlorite method (Lewis and Fleming 1995).

### RNAi

The RNAi treatment was carried out as described previously (Kamath and Ahringer 2003). The RNAi construct for *unc-22* (IV-6K06) was obtained from the RNAi library from the Medical Research Council. Synchronized L1 larvae were grown on RNAi bacteria until they were adults.

### Ftir-Based Imaging Method (FIM)

In this study we applied FIM developed to image Drosophila larvae (Risse et al. 2013; Risse et al. 2014b) to record multiple worms. In this system FTIR is used to illuminate the worms with infrared light. Given the appropriate optical properties (i.e. refractive indices) of acrylic glass and air and adequate angles of incidence the infrared light, which is directed into an acryl glass plate, is completely reflected at the interface of the acryl glass and air. Due to a refractive index of agar which is between the indices of acrylic glass and air, light with appropriate angles of incidence can enter and propagate within the agar layer. If worms are in contact with agar the total reflection of light is frustrated resulting in light with angles below the critical angles of the agar/acrylic glass/air interfaces. Thus, light scattered by the worms passes through the agar and the acrylic glass and is detected by a camera mounted beneath the glass (Figure 1A; (Risse et al.2013)). The use of FTIR generates high contrast images in which only the contact surfaces between worm and agar are visible in the recordings. The light intensity is controlled by pulse width modulation (PWM) of the infrared LEDs via a micro controller (Arduino 2560). For image acquisition a 4 megapixel camera (Basler Ace2040) was used equipped with a 7× macro zoom COMPUTAR TEC-V7X lens and an infrared long pass filter with a cut-off wavelength of 825 nm (Schneider Kreuznach IF093).

**Figure 1.**
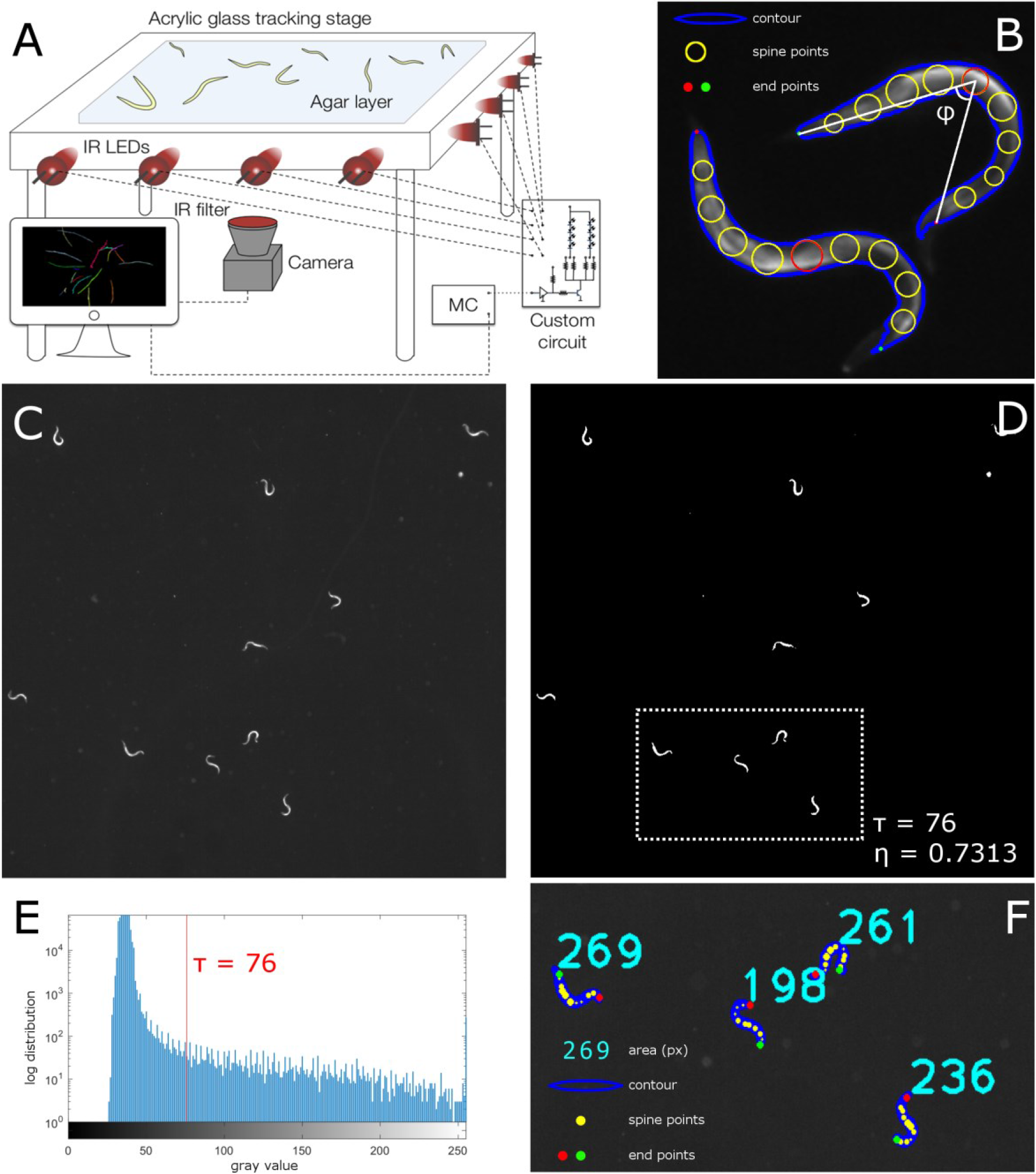
The FIM setup for *C. elegans*. **(A)** Schematic illustration of the FIM setup. IR = infrared light; MC = micro controller for LEDs (see (Risse et al. 2013) for detailed explanations). **(B)** Representation of worms in FIMTrack. The contour (blue line) and 11 spine points (yellow circle) including the head (red dot) and tail (green dot) of the worms are shown. The bending angle φ was determined between the end points (head and tail) and middle spine point 6 (red circle). **(C)** Raw image of *C. elegans* captured with FIM. Image was captured using a 4 megapixel Basler Ace2040 camera with a 7× macro zoom COMPUTAR TEC-V7× objective. **(D)** Binary image after applying Otsu’s thresholding. The overall optimal threshold τ=76 leads to a separability of η=0.7313 (where zero would indicate an image with pixels of only a single constant gray value and 1 would indicate a picture containing pixels of just two values; i.e. perfect separability (Otsu 1975)). **(E)** Histogram for the raw image shown in **(C)** (threshold τ indicated by a red bar). **(F)** Segmentation result using Otsu’s threshold (clipping from the binary image **(D)**; dashed box). The area of each detected contour is given in bright blue, the contours are highlighted in dark blue, spine points are given in yellow and the end points are drawn in red and blue.

### FIMTrack

FIMTrack is a freely available software (fim.uni-muenster.de) and computes posture and motion related features based on FIM images over time (Risse et al. 2013; Risse et al. 2014b). The worm posture is delineated by a contour detection algorithm, the spine is calculated and the head is identified as the point with the highest curvature (Risse et al. 2014b). The number of equidistant spine points can be manually defined. In this work 11 spine points were used, where spine point 1 displayed the head position, spine point 6 marked the centre of the animal and spine point 11 defined the tail. The centre of the worm was used to calculate the velocity as well as the travelled distance. Spine points 1, 6 and 11 were used to extract the bending angle φ (φ=180° is defined as no bending; i.e. the angle derived by the head, mid and tail point), which can be used to extract the thrashing rates of the worms as a changing rate from left bended (φ<180°) to right bended (φ>180°) (Figure 1B). During this work several extensions were implemented in the FIMTrack software. A “region of interest” (ROI) was added to restrict the analysis to a specific region within the image. Furthermore landmark areas could be set during the analysis to determine the localization of the chemical attractant/repellent (Figure 2B). If landmarks are used, FIMTrack computes the distance between worms and landmark, the bearing angle relative to the landmark and a binary indicator which indicates if the central spine point of an animal is within a landmark area or not. Additionally, tracks can be filtered by length via a manually set cut-off frame value. The subsequent statistical analysis of the obtained data was performed with customised MATLAB^®^ scripts and SigmaPlot^®^.

**Figure 2.**
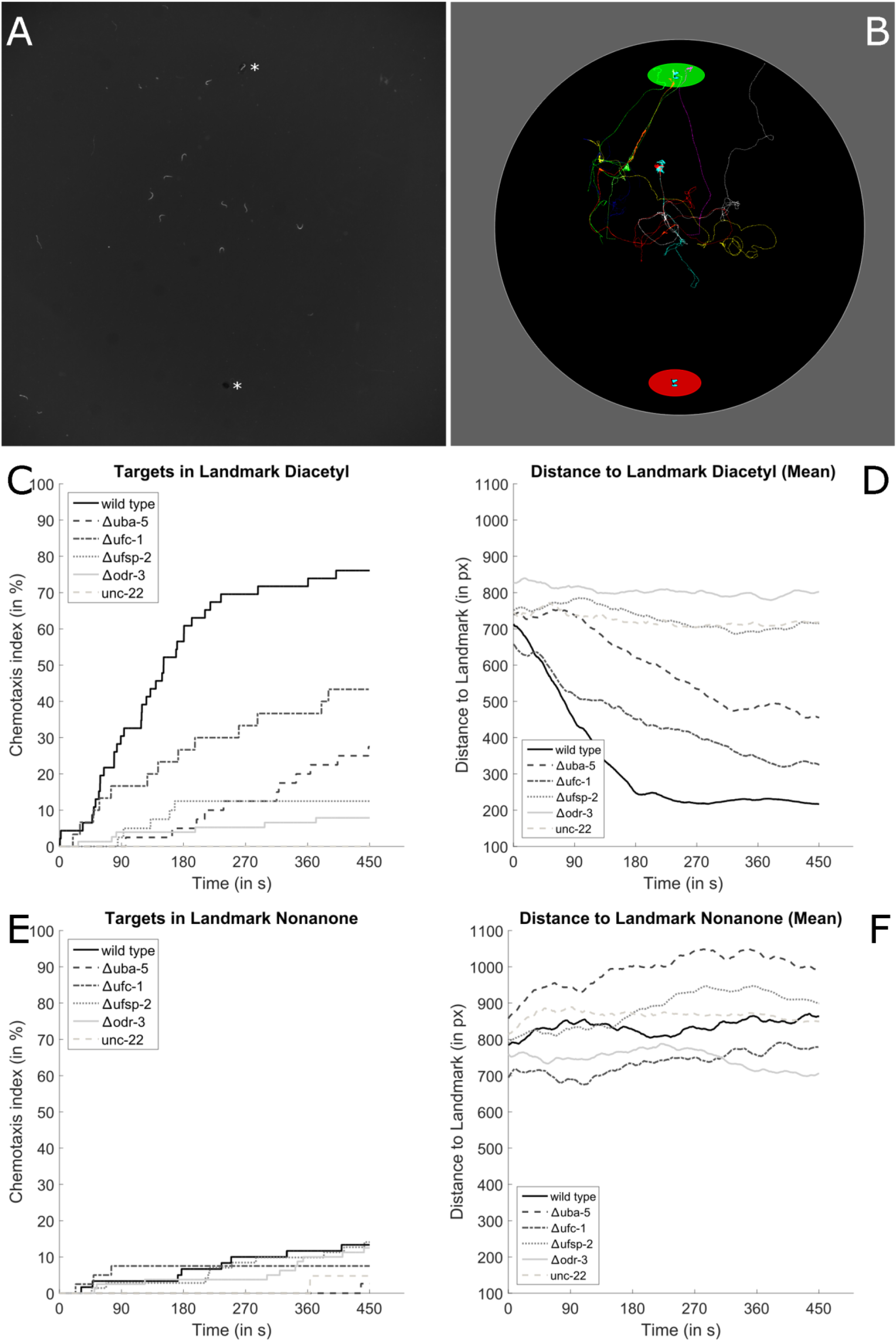
Chemotaxis. **(A)** FIM image of chemotaxis experiment. Small holes mark the landmarks as described in the methods and are tagged with asterisks. 10 synchronized worms were placed at the centre of the experimental plate. **(B)** Trajectories of *C. elegans* and landmark regions in the chemotaxis assay shown in **(A)** generated by FIMTrack. Worms were recorded for 450 s with 10 fps. Black circle = Region of Interest (ROI); green circle = attractant/repellent landmark A; red circle = ethanol control landmark B; coloured lines = trajectories of each worm. **(C)** Chemotaxis indices (CI) over time of the wild type (n=46), Δ*uba-5* (n=40), Δ*ufc-1* (n=30), Δ*ufsp-2* (n=40), Δ*odr-3* (n=76) and *unc-22* RNAi worms (n=15) using diacetyl at landmark A. **(D)** FIMTrack continuously measures the distance of each worm to landmark A. The mean distance of each strain towards diacetyl was determined at each time point and plotted using customised MATLAB^®^ scripts. **(E)** Chemotaxis index of the wild type (n=60), Δ*uba-5* (n=38), Δ*ufc-1*(n=40), Δ*ufsp-2* (n=71), Δ*odr-3* (n=80) and *unc-22* RNAi worms (n=21) using 2-nonanone at landmark A. **(F)** The mean distance of each strain towards 2-nonane was determined at each time point and plotted using customised MATLAB^®^ scripts.

### Chemotaxis

The chemotaxis assay is an extension of the method by Bargmann (Bargmann et al. 1993). The recording agar plates were prepared with an agarose concentration of 6 g/l. No other chemical additives were added and the agar was poured as thin as possible to create optimal transparency. Small holes were punctured in the agar to mark the two landmark areas approximately 6 cm apart on the same agar plate (Ø9 cm), because liquids are not visible in the FIM setup (Figure 2A). On one landmark a 1.5 μl drop of 1:1000 diluted diacetyl or 2-nonanone was placed and 1 μl of 0.5 M sodium azide was added. At the second landmark 1.5 μl of 99.9% ethanol was placed together with 1 μl of 0.5 M sodium azide. Ten synchronized adult worms were transferred to a bacteria-free NGM plate for 15 minutes. After this habituation phase the worms were located at the centre of the experimental plates. Trajectories of the worms were recorded for 450 s with 10 fps using FIM. The chemotaxis index was calculated using the FIMTrack and MATLAB^®^ software and is defined by the ratio between worms which central spine point is inside the set landmark region against the total number of worms over time. Furthermore, we used the time-coupled data of the chemotaxis index to analyse the phenotypic difference between the genotypes with the Kaplan-Meier (log rank) test with the Holm-Sidak method in SigmaPlot^®^. In addition, customised MATLAB^®^ scripts were used for subsequent analysis of the worm distance towards the landmarks with the Wilcoxon rank sum test.

### Thrashing

Thrashing is defined as the lateral swimming movement in which the worm body bends from one C-shape to the opposite ɔ-shape and vice versa (Buckingham and Sattelle 2009). The recording agar plates were prepared with an agarose concentration of 6 g/l. No other chemical additives were added and the agar was poured as thin as possible. 1.5 μl droplets of M9 were evenly distributed on the agar plate. Single adult worms were directly transferred from *E. coli* OP50 seeded NGM plates into each M9 droplet. The thrashing movement was recorded in at least three technical replicates for 60 s with 20 fps using the FIM setup. Image data was analysed using FIMTrack and customised MATLAB^®^ scripts were used to count the thrashing rate. A single thrash is defined by a change in the bending angle φ, which is defined between the spine points 1, 6 and 11, going from left bended (φ<180°) to right bended (φ>180°) and back again or vice versa.

### Electrotaxis

For the electrotaxis assay the recording agar plates were prepared with an agarose concentration of 10 g/l and 2 mg/l NaCl and 1850 μl glycerol were added to achieve electric conductivity and the agar was poured as thin as possible to create optimal transparency of the plate. Synchronized worms were placed on the recording plate. The custom made electrotaxis dome (Figure 4A) was placed on top of the FIM setup. Thereby two platinum wires got into contact with the agar layer and 60 V were applied, using an electrophoresis power supply (GE Healthcare EPS 601). The worms were tracked for 180 s at 10 fps in four technical replicates.

### Imaging of GFP Expression

For the GFP experiments the recording agar plates were prepared in the same way as for chemotaxis experiments. Synchronized wild type worms were placed together with worms of the GFP expression strain P*vit5*::GFP at the centre of the agar plate. The GFP experiments were conducted as described previously (Risse et al. 2016). Images were captured using two 4 megapixel Basler Ace2040 cameras with a Kowa LM35HC3 objective with a 5mm spacer ring. Overlay images were then computed as described in (Risse et al. 2016). Worms were tracked at 10 fps.

## Results

In this study we advanced the FIM technology to a novel multi-purpose worm tracking suite which has the ability to analyse movement patterns of multiple worms in various experimental approaches in detail. To verify the reliability of FIMTrack we tracked *C. elegans* worms on different agar concentrations as well as LED intensities. Verification was done by overlaying the videos with the tracking results and manually inspecting different features plotted in FIMTrack. An agar concentration of 6 g/L and LED intensity set to a PWM of 100% was chosen as optimal parameters. With this setting the FIM setup is capable of imaging and recognising *C. elegans* worms in detailed close ups as well as in 9×9 cm wide arenas (Figure 1). The applied thresholding is able to display a high separability value for all different experiments and resolutions. To quantify binary separability into foreground (i.e. worms) and background Otsu’s thresholding quality measure η can be utilised. A η=0 indicates an image with pixels of only a single constant grey value and η=1 indicates a picture containing pixels of two values; i.e. perfect separability (Otsu 1975). Given a FIM imaged *C. elegans* the overall optimal threshold τ=76 leads to a separability of η=0.7313, confirming that this imaging technique maximizes the contrast between foreground and background and that it is sufficient to extract the worms via thresholding (Figure 1D+E). Since FIMTrack is not yet able to resolve collision events between worms we reduced the number of such events by decreasing the number of simultaneously tracked worms to 10-32, depending on the experiment. In the following we demonstrate the capability of the novel worm tracker setup to reliably detect phenotypic differences between *C. elegans* strains in multiple behavioural experiments.

### Chemotaxis

Worm trackers are often used to analyse chemosensory phenotypes (Swierczek et al. 2011). We combined the theoretical setting of the chemotaxis assay of Bargmann (Bargmann et al. 1993; Dwyer et al. 1998) with the technical power of the FIM setup and FIMTrack. During the experiment each worm was continuously tracked for 450 s at 10 fps (Figure 2A). We have integrated a landmark implementation in FIMTrack to quantify the behaviour relative to these markers automatically. The landmark allows defining the area covered by the attractant/repellent and control chemicals (Figure 2B). With this additional information FIMTrack continuously measures the worm’s distances to the landmark during the whole experiment. Additionally, the chemotaxis index (CI) is calculated as the ratio between worms in the given landmark against the total number of worms over time (Figure 2C+E). Thus, the Kaplan-Meier test can be used to statistically compare the worms in the landmark over time between genotypes.

To test the functionality of the chemotaxis assay we used wild type worms and Ufm1 cascade mutant strains Δ*uba-5*, Δ*ufc-1* and Δ*ufsp-2* together with the chemosensory deficient mutant Δ*odr-3* (Roayaie et al. 1998) and movement impaired *unc-22* RNAi worms (Benian et al. 1989) (Figure 2C). In every movie 10 worms of each genotype were recorded for 450 seconds. Significant differences towards the wild type were detected by comparing the chemotaxis indices over time with the Kaplan-Meier test as described in the material and methods. As expected wild type worms displayed strong chemotaxis towards diacetyl, leading to a CI at the end of the experiment (t_450_) of 78% (n=46), whereas only 8% of the chemosensory deficient mutant *Δodr-3* mutant worms reached the diacetyl landmark after 450s (n=76). When the CI of *Δodr-3* is compared to wild type over time and statistically analysed with the Kaplan Meier test a significant difference is revealed (p<0.001×10^−7^). Furthermore, none of the movement impaired *unc-22* RNAi worms reached the diacetyl landmark (CI=0%; n=15; p=0.685×10^−3^). Therefore the baseline of reaching a landmark by random crawling was conservatively set to 10%. Interestingly, Ufm1 cascade mutants showed divergent results. As reported before (Chen et al. 2014) Δ*ufsp-2* worms display a similar CI compared to the chemosensory deficient mutant strain Δ*odr-3* (CI=12,5%; n=40; p=0.785×10^−7^). However, the Ufm1 cascade mutant Δ*ufc-1* had an intermediate CI of 43% (n=40; p=0.149), which is nonsignificant compared to wild type, but display significant differences to Δ*odr-3* (p=0.397×10^−5^,), Δ*ufsp-2* (p=0.00653,) and *unc-22* RNAi worms (p=0.0479). The Δ*uba-5* strain showed a further decreased CI of 27.5% (n=30; p=0.918×10^−6^), but no significant differences to the chemosensory deficient controls could be determined.

The capability of FIMTrack to fully conserve trajectories allowed calculating the mean distance of all worms from a landmark over time and the mean difference (MD) of the distance to the landmark from t_0_ to t_450_ could be used as an additional measure of chemosensory behaviour (Figure 2D+F). Wild type worms display a consistent approach towards the diacetyl landmark (MD=−492.2px). A similar behaviour was shown for Δ*ufc-1* (MD=−332.6px) and Δ*uba-5* worms (MD=−285.1px) although the slope of the function is smaller (Figure 2D). The chemosensory control Δ*odr-3* (MD=−24,8px), the movement impaired *unc-22* RNAi worms (MD=−26.6px) as well as Ufm1 mutant strain Δ*ufsp-2* (MD=−36.1px) only slightly alter their mean distance towards the diacetyl landmark over the course of the experiment. To distinguish the worm strains statistically, the area below the distance to landmark functions of each single worm was calculated and a Wilcoxon rank sum test was applied to the data. Compared to the chemosensory deficient mutant strain *Δodr-3* (n=76) a significant reduction of the distance towards the diacetyl landmark over time was shown for wild type (n=46, p=0.122×10^−10^), Δ*ufc-1* (n=40, p=0.387×10^−7^) and Δ*uba-5* worms (n=30, p=0.0014). Furthermore compared to wild type worms a significant difference of Δ*ufc-1* (n=40; p=0.045) and Δ*uba-5* (n=30; p=0.121×10^−6^) was shown, thus confirming the intermediate phenotype of these Ufm1 mutants.

In a second chemotaxis experiment the behaviour towards 2-nonanone, which is described as a repellent for *C. elegans* (Bargmann et al. 1993), was analysed for all previously described worm strains. All worm strains had a chemotaxis index near the baseline of randomly reaching a landmark of ~10% towards 2-nonanone (Figure 2E). Additionally, the Kaplan-Meier test did not indicate significant differences between the strains (p=0.445, n=300). The obtained mean distance functions of the strains showed that the wild type (MD=+79.1px, n=60), Δ*ufc-1* (MD=+85.1px, n=40) and Δ*uba-5* (MD=+113.1px, n=38) worms increased their distance to the 2-nonanone landmark over time (Figure 2F). Whereas the chemosensory deficient mutant Δ*odr-3* worms slightly decreased the distance (MD=−53.4px, n=80). The Wilcoxon rank sum test only detect a significant difference between Δ*uba-5* worms towards the control strain Δ*odr-3* (p=0.0011) as well as Δ*ufc-1* (p=0.0024).

Taken together FIMTrack is able to detect strong chemosensory phenotypes as well as subtle differences in Ufm1 mutants that were undetectable using a classical chemotaxis index due to its high data quality and trajectories which are consistent over time.

### Thrashing

Quantification of *C. elegans* swimming movement, called thrashing, is a widely used assay for measuring general mobility of *C. elegans* worms (Buckingham and Sattelle 2009). Thrashing assays are very time consuming and error-prone due to subjectivity when done manually. Only a single worm can be monitored at a time by human observation. Furthermore, automatic thrashing calculation can be aggravated due to light reflections of the water droplets if direct illumination is used. The use of FIM imaging circumvents this problem, because light is not scattered by the water the imaging is not disturbed. Instead it is even improving the sensitivity of this particular recording technique, since light is dispersed in the droplet and worms are illuminated from all sites (Figure 3A). We tested the ability of the FIM setup to display and automatically analyse thrashing behaviour of multiple worms at the same time. To investigate the accuracy of the tracking results, we manually counted thrashes in eight experiments with at least 10 worms each and compared the results with the automatically generated FIMTrack results. The automated analysis detected 8019 thrashes compared to two experimenters, who counted 8018 and 8098 thrashes, respectively. The variance between the two human countings and FIMTrack is less than 0.5%, therefore confirming the reliability of the automated analysis. Wild type worms displayed a median thrashing count of 159 thrashes/min (n=94), whereas *unc-22* RNAi treated worms exhibit a reduced median count of 118 thrashes/min (n=101; Figure 3B). The Ufm1 mutant Δ*uba-5* and Δ*ufc-1* as well as Δ*odr-3* worms display no significant difference to the wild type motility (one-way ANOVA, p>0.05). Interestingly, the one-way ANOVA analysis confirmed a significant difference between *unc-22* RNAi worms and all other strains (p<0.001), except for the Δ*ufsp-2* strain. Taken together the results confirm that FIM and FIMTrack is a suitable and accurate tool to analyse thrashing phenotypes.

**Figure 3.**
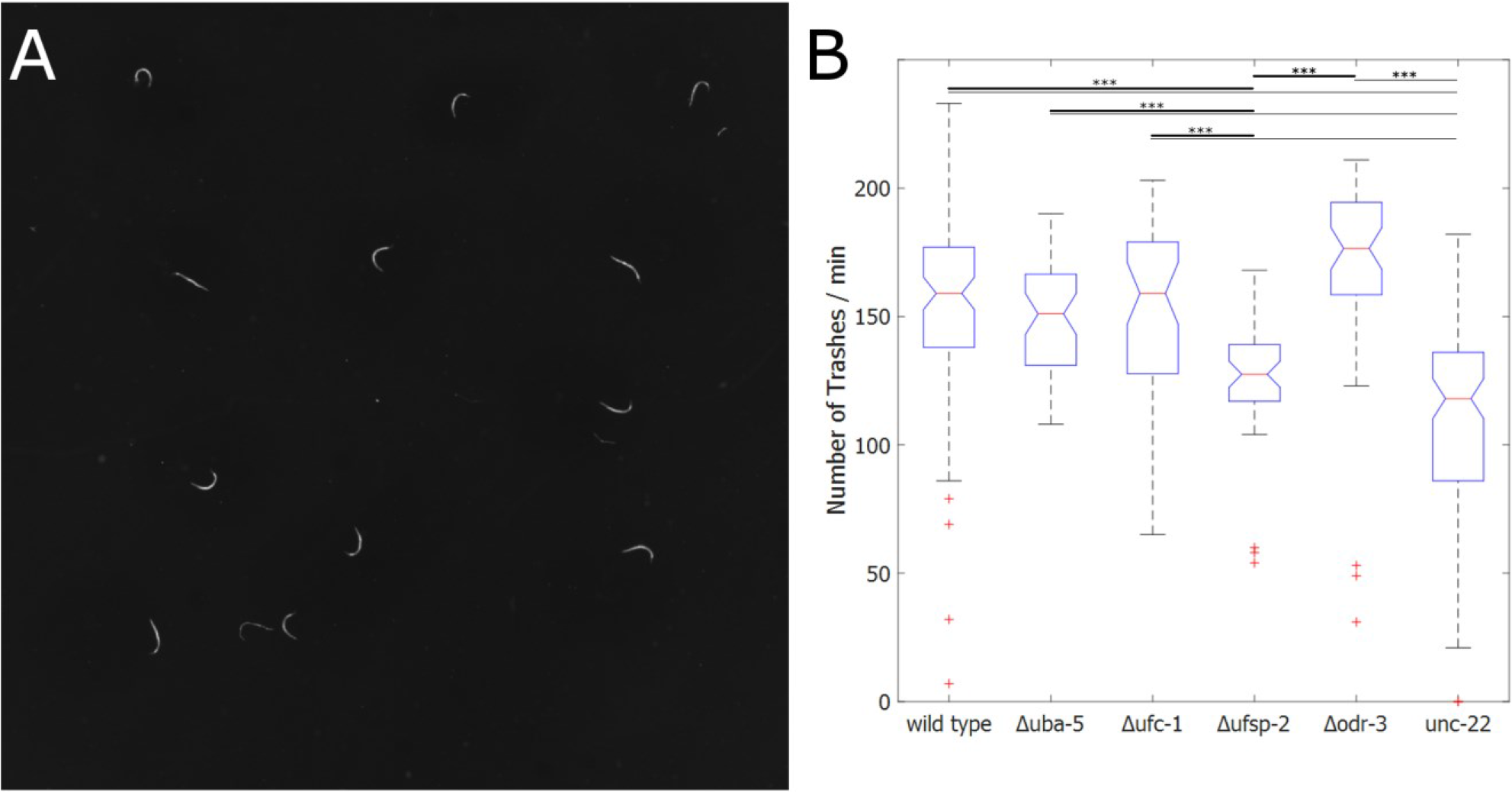
Thrashing. **(A)** FIM image of thrashing experiments. Each worm is placed in a 1.5μl droplet of M9 medium. M9 droplets are not visible in FIM images. **(B)** Boxplot of thrashing count of wild type (n=94), Δ*uba-5* (n=48), Δ*ufc-1* (n=45), Δ*ufsp-2* (n=46), Δ*odr-3* (n=48) and *unc-22* RNAi worms (n=101). One thrash was counted as a change in the bending angle φ going from left bended (φ<180°) to right bended (φ>180°) and back again or vice versa. The median number of trashes per minute is displayed as a red line. One-way ANOVA was calculated with MATLAB®. Thick lines represent significant differences towards Δ*ufsp-2* whereas thin lines mark significant differences towards *unc-22* RNAi worms and are marked with asterisk (***: p<0.001).

### Electrotaxis

Another behavioural parameter to differentiate genotypes is the electrotaxis of *C. elegans.* Worms move at their maximum speed to the negative pole in electric fields with an inclination that rises with the strength of the field (Gabel et al. 2007). The electrotaxis behaviour differs in movement patterns and speed, without being corrupted by inconsistent kinetics occurring in free movement experiments (Maniere et al. 2011). Based on Maniere et al., we developed a straightforward and easy-to-use electrotaxis dome for the FIM setup (Figure 4A). With this extension the generation of an electric field on the recording agarose plate using a common electrophoresis power supply was possible. It is noteworthy, that this setup would not be usable in an environment with direct illumination. We tested this setup with wild type and *unc-22* RNAi treated worms. We imaged the worms for 180 s with 10 fps at 60 V. Most of the worms displayed continuous movement towards the negative pole with varying inclination (Figure 4B). Immobile worms were automatically excluded. The wild type worms moved with an average speed of 152 μm/s while *unc-22* RNAi worms moved with 122 μm/s. A two-tailed t-test revealed a highly significant difference between the two groups (p<0.001) (Figure 4C). Therefore the FIM based worm tracker can be used efficiently to analyse the mobility of *C. elegans* in electrotaxis behaviour.

**Figure 4.**
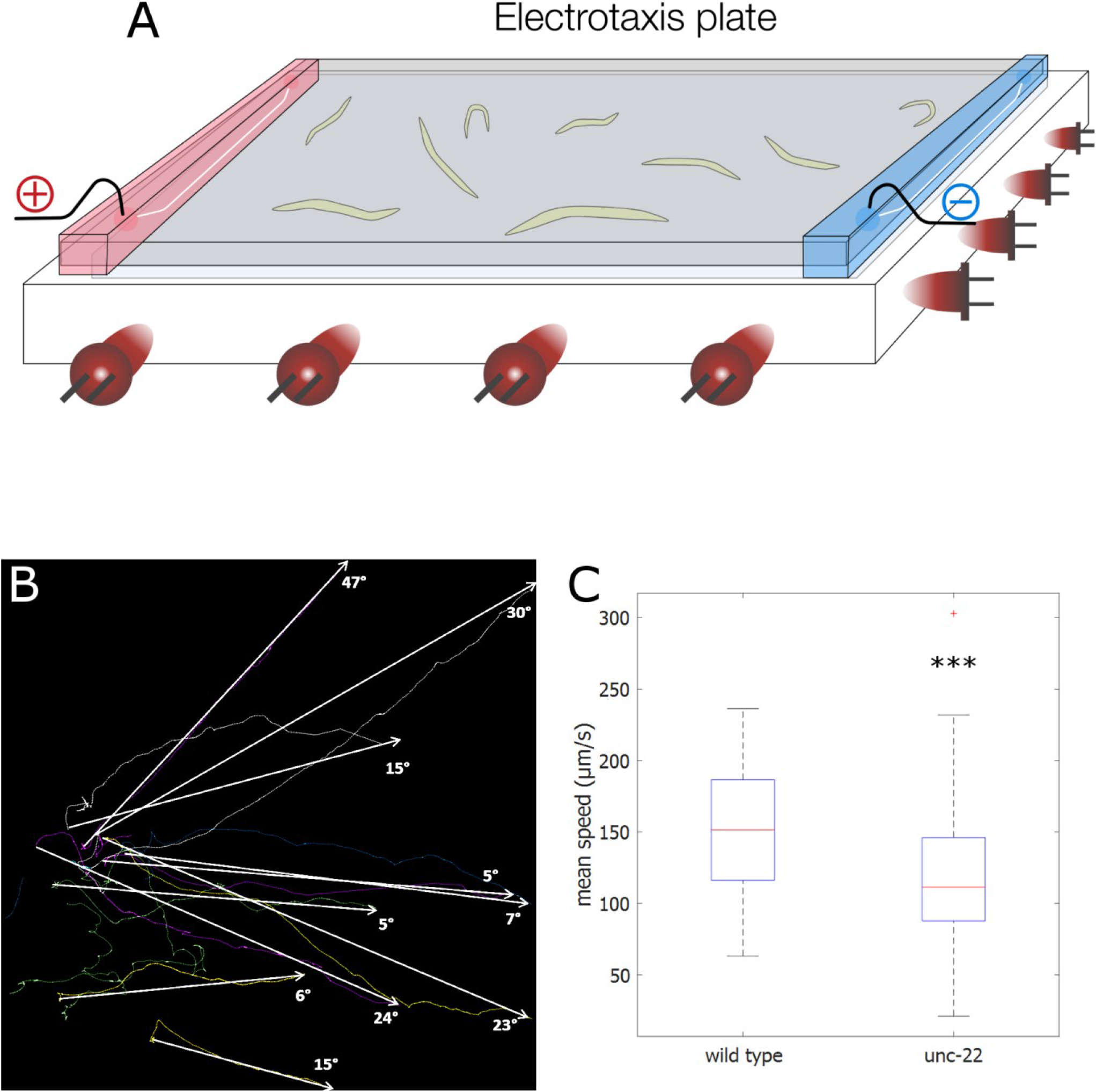
Electrotaxis. **(A)** Scheme of the electrotaxis dome. For details see materials and methods. **(B)** Example of trajectories of 10 wild type worms in electrotaxis experiment moving towards the negative pole recorded for 180 s at 10 fps. Angles of inclinations between start and end time point are shown with arrows. **(C)** Mean speed of wild type (n=47) and *unc-22* RNAi worms (n=55). Significant differences are marked with asterisk (two-tailed t-test, ***: p<0.001).

### GFP Imaging

In addition, we analysed whether the FIM setup is also able to detect GFP signals in *C. elegans* worms. It has already been shown for *Drosophila* larvae (Risse et al. 2013) and has recently been integrated further into the multi-colour, multi-purpose imaging system FIM^2C^ to incorporate optogenetic tools as well as to resolve collision events of Drosophila larvae (Risse et al. 2016). Here we show that we were able to apply this technique in principle to the much smaller *C. elegans* worms, demonstrated by the capability of FIMTrack to distinguish GFP-labelled P*vit5*::GFP worms from non-labelled wild type worms (Figure 5). Nevertheless, further work needs to be done to incorporate tissue-specific expression of GFP and optogenetics in the more size-challenging *C. elegans* worms. However, the use of labelled and non-labelled GFP worms already creates the possibility of analysing two genotypes on the same plate, which facilitates the prospect for further behaviour analyses.

**Figure 5.**
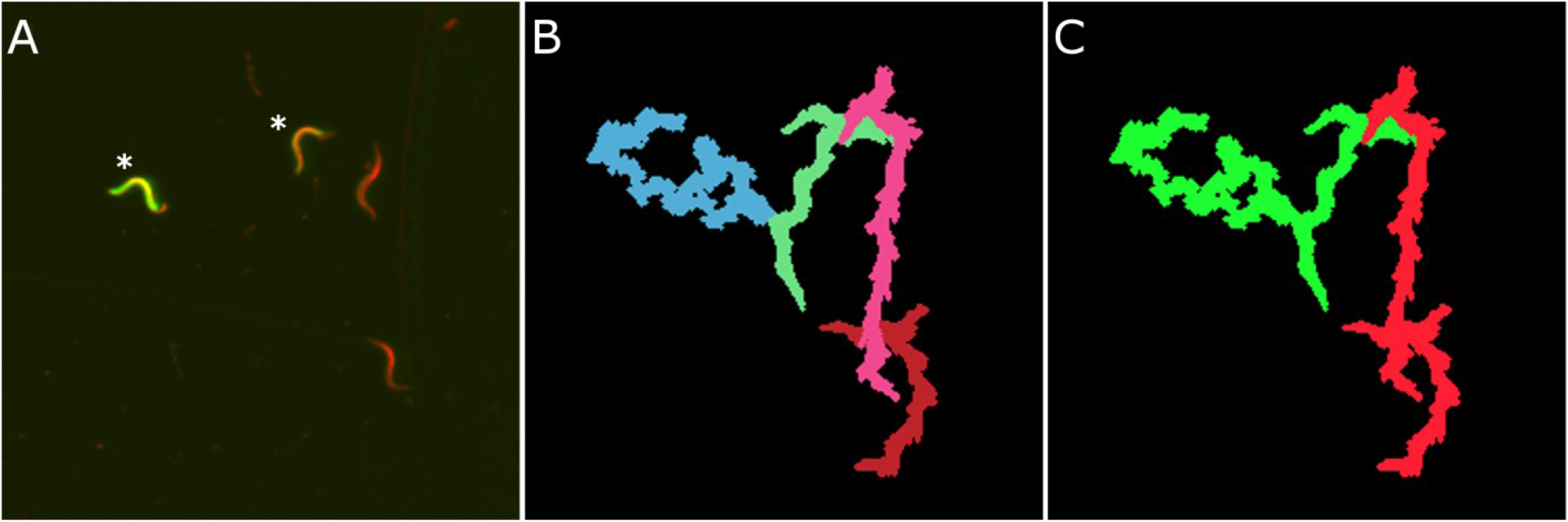
GFP imaging applying FIM^2C^. **(A)** Overlay FIM image of GFP experiments. GFP expressing P*vit5*::GFP worms are marked with asterisks. **(B)** Trajectories of worms displayed in (A) generated with IR image information by FIMTrack. **(C)** Trajectories of GFP-labelled worms (green) and non-GFP worms (red) are segmented due to additional information (Risse et al. 2016).

## Discussion

The identification of behavioural phenotypes under different conditions is crucial for the general understanding of motor neuronal functions, participating cellular pathways and single proteins in *C. elegans.* Numerous studies demonstrated the usefulness of worm trackers in such analyses (Brown and Schafer 2013; Krajacic et al. 2012; Polli et al. 2015; Ramot et al. 2008; Restif and Metaxas 2008). However most of the used worm trackers are tackling a specific behavioural phenotype, track only single worms at a time or use a centroid-based algorithm. The most commonly used “Multi Worm Tracker” overcomes these limitations (Swierczek et al. 2011), but rejects all colliding worms and does not save the image data by default for post-processing, leading to a possible loss of informative data. Furthermore, most trackers have to cope with illumination and contrast disadvantages arising from bright field image illumination. Here we described a novel worm tracker which utilises FIM as a basis leading to high contrast images of small and translucent *C. elegans* (Figure 1). The imaging quality is not impaired by the moist surface of agar, liquids placed on the agar or even devices covering the tracking arena. On the contrary, liquids have a neutral or even beneficial effect on imaging and the possibility of placing additional devices on top of the tracking arena generates new opportunities for behaviour experiments like electrotaxis (Figure 4) or thermotaxis (Risse et al. 2014b).

Besides the imaging advantage of the new setup, its accuracy, sensitivity and time-efficiency are also crucial advantages for screening approaches. During the chemotaxis experiment this worm tracking method continuously calculates the chemotaxis index over a short time (450s), whereas former studies took 60 or 90 minutes as experimental time and measured the chemotaxis index at different intervals (Chen et al. 2014; Matsuura et al. 2013; Bargmann et al. 1993). Furthermore, our analysis showed that more than 50% of the wild type worms covered a ~ 2 cm distance to the diacetyl landmark in less than three minutes. This result underlines the importance of short, highly monitored chemotaxis assays to identify slight aberrations from wild type behaviour, which might be fading out over a longer time span. Concordantly, our data showed that the novel tracking setup is able to distinguish intermediate chemosensory phenotypes of Ufm-1 cascade mutants Δ*uba-5* and Δ*ufc-1* (Figure 2). Both strains display a reduced CI and a slower distance reduction towards diacetyl compared to wild type worms. However a tendency towards diacetyl was shown, which might mask this intermediate phenotype using the classical chemotaxis index as it was used before (Chen et al. 2014).

In a next step, we successfully adopted the FIM setup for quantification of *C. elegans* swimming movement, which is a suitable assay to detect motility defects. Several automated worm trackers are specialized on this analysis and use covariance matrices (Buckingham and Sattelle 2009), curvature contour plots (Krajacic et al. 2012) or a vector based model (Restif and Metaxas 2008). FIMTrack on the other hand uses the spine points at the head, centre and tail of the measured contour to extract the bending angle φ (Figure 1B). This straight-forward approach displays a high accuracy in thrashing count and confirmed the impaired movement ability of *unc-22* RNAi treated worms, whereas the Ufm1 mutant strains Δ*uba-5* and Δ*ufc-1* as well as Δ*odr-3* displayed a wild type phenotype (Figure 4). In combination with the chemotaxis result it is thus possible to distinguish an impaired movement phenotype (*unc-22* RNAi) from an impaired chemosensory phenotype (Δ*odr-3*). Thus, confirming that the intermediate phenotype of Δ*uba-5* and Δ*ufc-1* worms in the chemotaxis experiment is solely dependent on a chemosensory defect. It is noteworthy, that Δ*ufsp-2* worms display a reduced motility and therefore the highly reduced chemotaxis index might be a combination of a chemosensory and a motility defect. This is the first time that a motility deficiency of Δ*ufsp-2* worms has been shown.

As a third behavioural assay we implemented an electrotaxis assay and measured the crawling speed of *C. elegans* in an electric field. The electrotaxis behaviour of the worms suppresses complex movement patterns like reversals and omega-bends and leads to crawling with maximum velocity towards the negative pole (Gabel et al. 2007). In our experiments we confirmed the previously detected impaired movement phenotype of *unc-22* worms. since the electrotaxis assay allows an easy and direct measurement of the speed and motility, it can be used to sort different phenotypes and screen for proteins or therapeutic biomolecules associated with muscle or neurodegenerative diseases (Maniere et al. 2011).

At last we show that the novel worm tracker is capable to detect and distinguish GFP-labelled worms from non-labelled worms (Figure 5). This might lead to a resolution of collision events which is the biggest loss of information in our setup as well as in all other multi-worm trackers. Wang et al. described an algorithm to improve tracking worms during omega bends and to some extend during collision events which combined locomotion state recognition and worm shape tracking (Yu Wang and Roysam 2010), but future work needed to be done to refine the results. Thus, no tracker is capable of efficiently solving collisions in a multi worm environment, yet. Most trackers only omit tracks of touching worms by using a size range and treat the trajectories before and after a collision as different worms or by deleting all incomplete tracks (Ramot et al. 2008). In consequence, many trajectories are not analysed leading to a loss of data and many incomplete tracks result in diminished data quality. The FIMTrack software also uses a size range of the area of the worms to distinguish between individual and colliding animals and omits colliding worms. However it features the possibility to manually merge worm trajectories in a quick and easy post processing step, making analysis over the whole course of the experiment possible. Nevertheless, an automatic algorithm to detect and resolve collision events would be favourable. It was shown recently that the FIM setup is able to resolve collision events of Drosophila larvae by the use of GFP (Risse et al. 2016). With the general ability to detect GFP in *C. elegans* as a basis, further experiments and adjustments will be conducted to transfer these results to the smaller *C. elegans* worms.

In conclusion, we adapted the FIM setup originally used to analyse larval locomotion of Drosophila to be applied to *C. elegans* with a broad range of applications like chemotaxis, thrashing, electrotaxis and GFP experiments. We confirmed the reliability and suitability of our setup in all experiments. Further applications like counting omega bends, reversals or thermotaxis can be easily implemented into the FIMTrack software if required. The FIM based worm tracker is a reliable high-throughput imaging and tracking system which provides a powerful and expandable tool box for multiple behaviour assays and therefore refine screening and phenotyping of *C. elegans* strains in various biological fields like investigation of neurodegenerative diseases, cellular pathways, stress cascades, chemo sensation and many more.

## Acknowledgement

We thank Prof. Dr. Christian Klämbt for his kind support and access to his facilities.

